# Microbiome impact of ibezapolstat and other *Clostridioides difficile* infection relevant antibiotics using humanized mice

**DOI:** 10.1101/2024.11.06.622322

**Authors:** Trenton M. Wolfe, Jinhee Jo, Nick V. Pinkham, Kevin W. Garey, Seth T. Walk

## Abstract

**Background:** Ibezapolstat (IBZ) is a competitive inhibitor of the bacterial Pol IIIC enzyme in clinical development for treatment of *Clostridioides difficile* infection (CDI). Previous studies demonstrated IBZ carries a favorable microbiome diversity profile compared to vancomycin (VAN). However, head-to-head comparisons with other CDI antibiotics have not been done. The purpose of this study was to compare microbiome changes associated with IBZ to other clinically used CDI antibiotics.

**Methods:** Groups of germ-free (GF) mice received a fecal microbiota transplant from one of two healthy human donors and were subsequently exposed to either IBZ, VAN, fidaxomicin (FDX), metronidazole (MET), or no antibiotic (control). 16S rRNA encoding gene sequencing of temporally collected stool samples was used to compare gut microbiome perturbation between treatment and no-drug control groups.

**Results:** Among the tested antibiotics, the most significant change in microbiome diversity was observed in MET-treated mice. Each antibiotic had a unique effect, but changes in alpha and beta diversity following FDX- and IBZ-treated groups were less pronounced compared to those observed in VAN-or MET-treated groups. By the end of therapy, both IBZ and FDZ increased the relative abundance of Bacteroidota (phylum), with IBZ additionally increasing the relative abundance of Actinomycetota (phylum).

**Conclusion:** In microbiome-humanized mice, IBZ and FDX had smaller effects on gut microbiome diversity compared to VAN and MET. Notable differences were observed between the microbiome of IBZ- and FDX-treated groups, which may allow for differentiation of these two antibiotics in future studies.

## Introduction

*Clostridioides difficile* is a Gram-positive, spore-forming, anaerobic bacterium and a leading cause of healthcare-associated and community infections ^1^. *C. difficile* infection (CDI) occurs when ingested spores germinate in the gut in response to changes in the gut microbiome induced primarily by antibiotics ^2^. Two major bacterial phyla, Bacillota and Bacteroidota, are typically the most negatively affected by common CDI-predisposing antibiotics, while the relative abundance of other phyla, like Pseudomonadota, increases ^3^. CDI is often treated with antibiotics, including vancomycin (VAN) or fidaxomicin (FDX). Metronidazole (MET) was previously a treatment of choice yet it is no longer recommended and is used only in certain settings due to decreased susceptibility against *C. difficile* ^4^. The ideal antibiotic for CDI treatment would selectively target *C. difficile* while preserving the rest of the microbiome, thus allowing for recovery of microbiome diversity and preventing CDI recurrence. Thus, development of new CDI antibiotics aims to minimize impacts on the microbiome. However, the distinct mechanisms of action of these drugs lead to varying effects on microbiome taxa. A direct way to evaluate a drug’s selectivity is to quantify changes in ecologic diversity (i.e., richness and evenness of microbiome taxa) between treated and untreated individuals, also known as beta diversity. FDX was approved for the treatment of CDI in 2011 and is now the prototypical narrow-spectrum antibiotic for CDI. This macrolide targets bacterial RNA polymerase (RNAP), which in turn inhibits transcription. A conserved residue in RNAP confers greater specificity to *C. difficile* and other Bacillota as opposed to Bacteroidota ^5^. This contrasts with the broad-spectrum activity of VAN, a glycopeptide that binds lipid-II-D-Ala-D-Ala to halt cell wall formation. VAN is active against both Bacteroidota and Bacillota ^6^. MET, a nitroimidazole prodrug, is activated within cells by oxidoreductases to produce reactive species that cause DNA and protein damage, depletion of thiols, and eventually lead to cell death ^7^. MET has been shown to reduce Bacillota but not Bacteroidota or Pseudomonadota ^8^.

Ibezapolstat (IBZ) is a novel antibiotic currently in clinical development for the treatment of CDI ^9^. IBZ acts as a competitive inhibitor of the C-family DNA polymerase IIIC (Pol IIIC) DNA synthesis substrate 2’-deoxyguanosine 5’-triphospate (dGTP) ^10^. The IBZ base pairing domain mimics guanine, resulting in the formation of an inactive ternary complex of IBZ, DNA, and PolC. The *pol IIIC* is an attractive antibiotic target as it is present in the phylum Bacillota, which includes *C. difficile*, and absent in other phyla common to the human gut (Actinomycetota, Bacteroidota, and Pseudomonadota) ^11^. In clinical trials, IBZ demonstrated targeted activity within the Bacillota phylum, ^9,12^,13 selectively killing *C. difficile* while maintaining or even increasing the relative abundance of closely related Pol IIIC Bacillota species known to be important in preventing recurrent CDI.

Despite the importance of microbiome disruption in the pathogenesis of CDI, a head-to-head comparison of all four CDI-relevant antibiotics on the same human microbiome has not been evaluated. Such a comparison is challenging in CDI patients due to inter-individual variability in microbiome composition and the diverse effects of CDI-predisposing antibiotics (i.e., before CDI onset). In this study, we treated two different groups of microbiome-humanized mice (i.e., germ-free recipients of human fecal transplants) with IBZ, FDX, VAN, MET, or untreated (no-drug) controls. While all antibiotics significantly impacted microbiome diversity, both IBZ and FDX resulted in significantly less perturbation compared to VAN or MET. These results are consistent with clinical trial results and demonstrate the value of humanized mice for preclinical evaluation of microbiome perturbation when developing prototypical therapeutics.

## Methods

### Animals

Animal experiments were conducted at Montana State University’s Animal Resource Center, an American Association for the Accreditation of Laboratory Animal Care (AAALAC)-accredited facility. All animal experiments were approved by the Montana State University Institutional Animal Care and Use Committee (IACUC). GF mice were housed in hermetically sealed and HEPA-filter ventilated vinyl isolators (Park Bio, Groveland, MA) and fed sterile food (5010 - Laboratory Autoclavable Rodent Diet, LabDiet, St. Louis, MO) and sterile water. All food and water supplied to GF mice were quarantined, monitored, and tested for microbial contamination via cultivation-dependent and cultivation-independent methods. For humanization, mice were given 100µL of thawed human stool mixed at 1:1 gram stool per volume of anaerobic PBS as previously described ^14^. Following humanization, mice were kept in a pre-sterilized biosafety cabinet. Due to the water insolubility of some of the antibiotics, clinically used antibiotic tablets were pulverized and mixed into a powdered mouse diet (AIN-93G Purified Rodent Diet, Dyets, Bethlehem, PA) and delivered in powder diet dishes (#910018 Complete Jar Set, Dyets, Bethlehem, PA)). Based on a previous study, ^15^ mice were acclimated twice; once to humanization (7 days) and subsequently to the powdered diet (7 days) prior to antibiotic administration to account for microbiome changes due to switch to powdered diet. Mice were housed under specific pathogen-free conditions (including murine norovirus) in individually ventilated, sterilized cages with food and water provided *ad libitum*. Mice were treated with antibiotics for 10 days, the guideline-recommended duration for CDI. A graphical illustration of the study design is provided in Figure 1.

**Figure 1.**
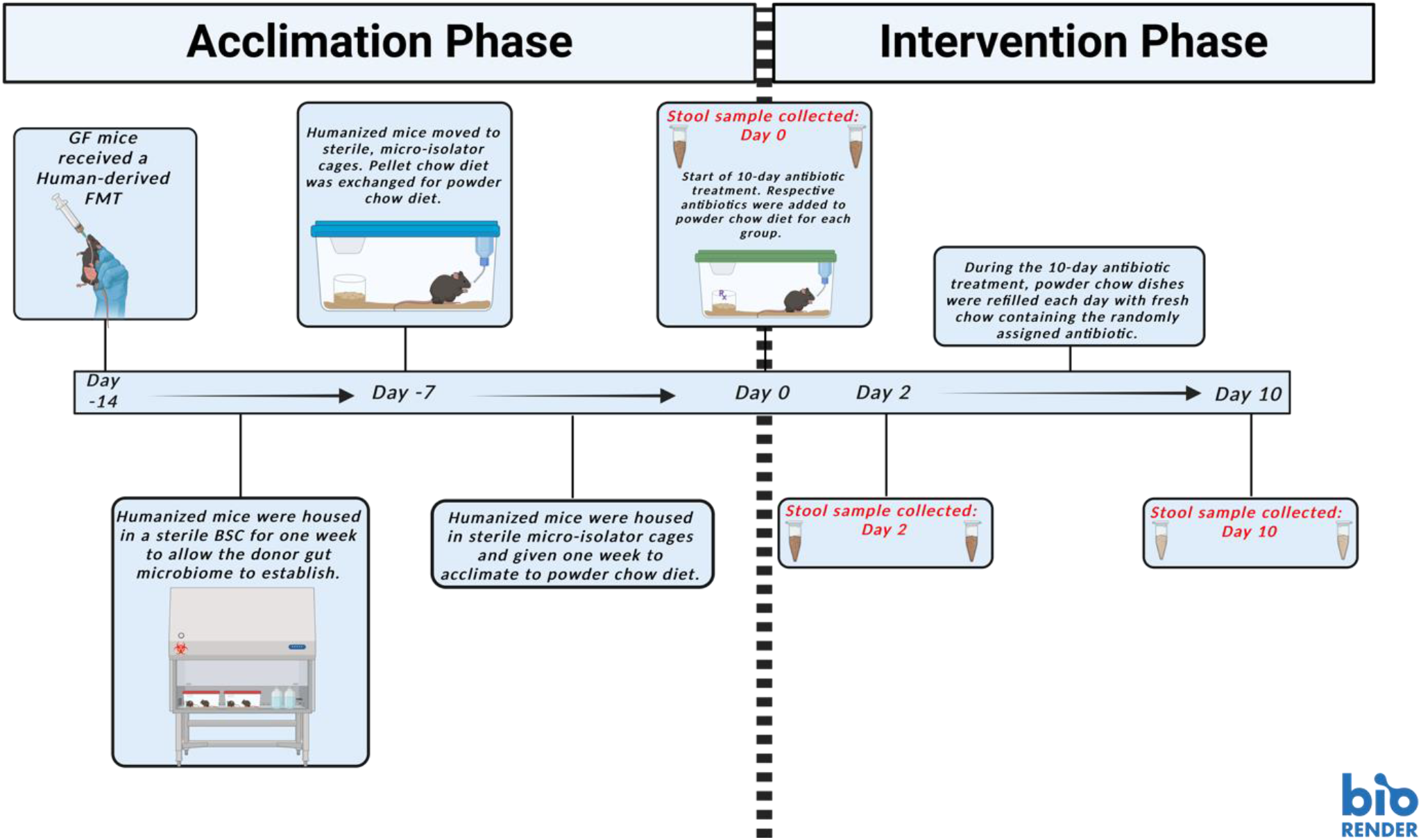
Graphical illustration of the humanized mouse model used in this study. Germ-free (GF) mice were engrafted with a healthy human donor microbiome via fecal microbiota transplant (FMT). Following FMT, mice were given seven days to establish a baseline microbiome. From there, conventional, pellet-style chow was exchanged for powder-style. Another seven days were given for natural and expected shift of the microbiome following this change in diet. At treatment day 0, a stool sample was collected to establish baseline microbiome. Mice were randomly assigned to an antibiotic-treatment group and stool samples were collected at treatment day 2 and 10. This figure was created with BioRender.com.

### Human Samples and Fecal Microbiota Transplantation (FMT)

Two frozen stool samples from healthy male adults (22 and 44 years of age) participating in a previous, IRB-approved study ^16^ were used to humanize mice via FMT in two trials. Samples were initially processed inside an anaerobic chamber and stored at −80°C until use. The frozen stool was thawed inside an anaerobic chamber (Coy Laboratories), suspended in sterile, pre-reduced phosphate-buffered saline (PBS) at an approximate stool weight to PBS volume ratio of 1:1, and briefly vortexed (<15 seconds) to generate a fecal slurry. 100 µL of this fecal slurry was used to inoculate sixteen GF mice in each trial via oral gavage (Supplemental Table 1).

### Pharmaceutical Drugs and Dosing

Human equivalent dosing of each drug was calculated based on US FDA “Guidance for Industry” ^17^. This allometric conversion adjusts drug doses in lab animals according to differences in body surface area between humans and mice. The same doses per gram body weight used to treat human CDI were converted to mouse equivalents (Supplemental Table 2). Antibiotic tablets were ground via mortar and pestle into a fine powder, weighed, and mixed with the powder diet. Mice were randomized into five treatments groups, corresponding to four CDI antibiotics (IBZ, FDX, VAN, MET) or an untreated (control) group for ten days per IDSA-SHEA CDI treatment guidelines ^4^.

Fresh powdered diet of diet/antibiotic mixtures were supplied daily. Powdered, pharmaceutical-grade antibiotics were added at the following concentrations: IBZ (45.34 mg/cage (n=5)/day in 40 g powdered chow/day), FDX (20.15 mg/cage (n=5)/day in 40 g powdered chow/day), VAN (25.19 mg/cage (n=5)/day in 40 g powdered chow/day), MET (50.37 mg/cage (n=5)/day in 40 g powdered chow/day). An untreated control group served as a comparator (40 g of powdered chow/day). A more detailed table of drug dosing is provided in Supplemental Table 2.

### 16S rRNA encoding gene sequencing

Unique ear punches identified individual mice and stool samples were collected seven days after acclimating to FMT, seven days after acclimating to the powdered diet (day 0 of treatment), day 2 of antibiotic treatment, and day 10 of antibiotic treatment. The stool was obtained by humanely restraining mice and collecting pellets directly into sterile Eppendorf tubes. Samples were immediately frozen at –80ºC until DNA extraction. Bulk DNA was extracted from thawed samples using the DNeasy® PowerSoil® Kit (Qiagen, Hilden, Germany). DNA was then frozen at –80ºC and shipped overnight on dry ice to the University of Michigan Center for Microbial Systems for Illumina MiSeq amplicon sequencing of the V4 variable region of the 16S rRNA encoding gene (dual-indexed barcoded primers, 2 × 250 base-pair reads) ^18^. Raw sequencing reads were processed using mothur (v.1.48.0)^19^ and the MiSeq Standard Operating Procedure (SOP) ^18^ accessed July 11, 2023 (https://mothur.org/wiki/miseq_sop/). Paired-end reads were assembled into contigs, screened for length and quality (maximum 275 bp and minimum of 246 bp with no ambiguous base calls), and aligned to coordinates 1968 through 11550 of the Silva ribosomal RNA gene reference database (v.138.1). Potential chimeras were identified and removed using the Uchime algorithm via mothur ^20^. Non-target reads (i.e., mitochondria, chloroplast, Eukaryota, and sequences unclassified at the domain level) were removed, and OTUs were identified in mothur using VSEARCH distance-based greedy clustering algorithm at the 97% sequence similarity threshold ^21^. From this, an OTU-based data matrix was built. Rare OTUs with <100 total reads were removed to minimize the influence of spurious observations. This resulted in the loss of one mouse in the IBZ-treated group in trial 2. Taxonomic classifications were assigned using the Bayesian classifier of the Ribosomal Database Project ^22^ implemented in mothur (training set v.18). Bray-Curtis (BC) distances between centroids were tabulated by first calculating the centroid of each group by averaging the abundance of each OTU across all communities in the group, and then taking the Bray-Curtis distance between the centroids.

### Statistical Analysis

Microbiome analysis was done with R version 4.3.1. Diversity estimates were calculated with R’s vegan package (version 2.6-4). Alpha diversity was quantified with the inverse of the Simpson index and compared using t-tests. Beta diversity was quantified using Bray-Curtis Dissimilarity, and PERMANOVA (using 999 permutations) was used to test for significance. Taxonomic changes were explored with Wilcox-ranked sum tests using the R package CAF’s (https://github.com/nvpinkham/CAF) taxonomy shared function. Statistical summaries are provided in Supplemental Data.

## Results

Two groups of GF mice (n=16 each) were evaluated in separate experiments (trial 1 and trial 2). Each trial consisted of fecal microbiota transplant (FMT) from a healthy human donor, with a different donor sample used in each trial. All treatments were well tolerated, except for one instance. In trial 2, one mouse in the MET treatment group was found dead on day 3. A necropsy did not reveal any gross anatomical abnormalities, and the cause of death remained unclear. No other mice exhibited noticeable signs of disease. Additionally, one mouse from the IBZ treatment group in trial 2 was excluded from analysis due to insufficient sequencing reads from stool samples. As expected, the microbiomes of humanized mice in trial 1 were significantly different from those in trial 2 before antibiotic treatment (Figure 2, PERMANOVA: R^2^=0.592, F_tat_=43.598 p=0.001), supporting that inter-individual differences in human donor microbiomes were successfully transferred to GF recipient mice via FMT. Antibiotic treatment effects were apparent by day 2 in both trials compared to untreated controls and persisted throughout the experiment until day 10 (Figure 2). Bray-Curtis dissimilarity beta-diversity indicated that microbiomes from IBZ- and FDX-treated mice were more similar to the no-drug control group compared to VAN- and MET-treated mice (Figure 3AB). By the end of treatment, VAN- and MET-treated mice showed significantly higher Bray-Curtis distance from the no-drug control (t-test, p<0.05) whereas IBZ and FDX-treated mice did not show significantly differences (Figure 3AB). Inverse Simpson’s index alpha diversity decreased in all antibiotic treatment group. However, IBZ did not show a significant difference from controls on day 2 or 10 of treatment (Figure 4). Alpha diversity of the FDX-treated group was significantly different from controls on day 2 or 10, while VAN and MET were significantly different on both days (Figure 4).

**Figure 2.**
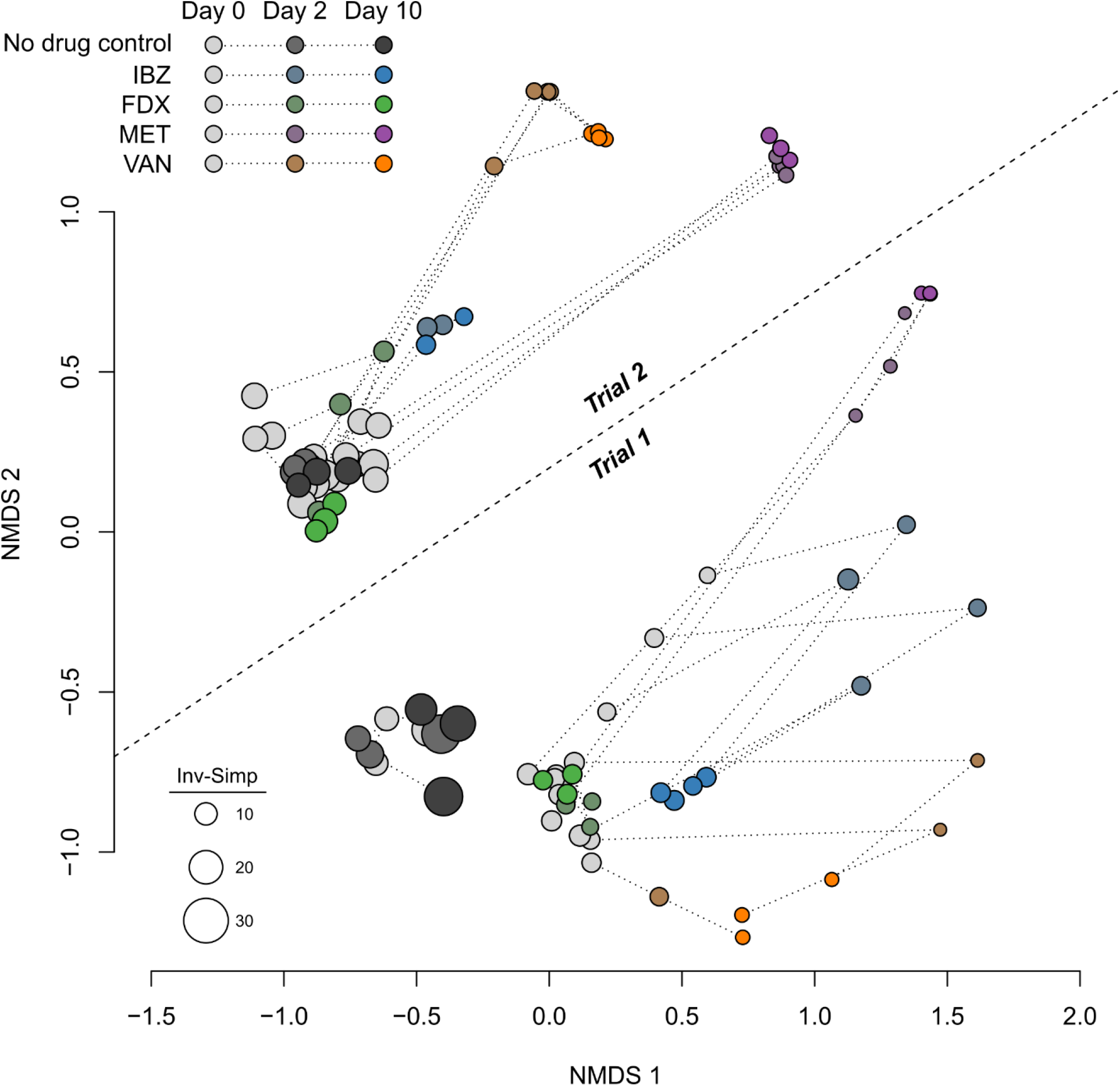
Non-metric multidimensional scaling (NMDS) of beta-diversity between microbiome samples. Baseline (day 0) samples are colored gray for all treatment groups. Days 2 and 10 of treatments are colored as shown in the legend. Dotted lines connect samples from individual mice. Trials 1 and 2 microbiomes were distinct as indicated by the labeled dashed line. The size of each dot was scaled according to alpha diversity (Inverse Simpson’s index), with larger dots representing more diverse microbiomes.

**Figure 3.**
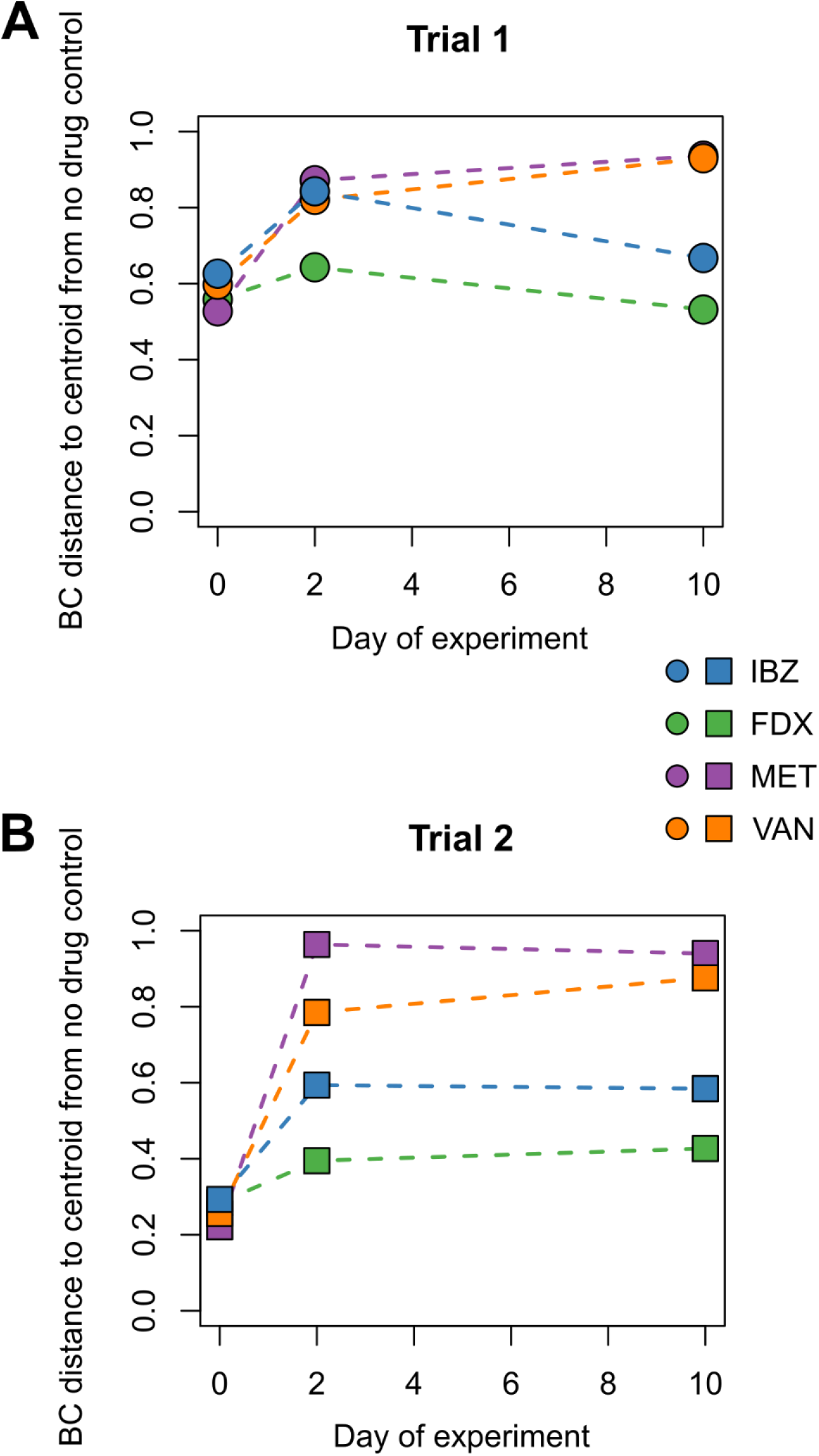
Bray-Curtis dissimilarity distance to treatment group centroids for each antibiotic relative to the no-drug control group. Trials 1 (A) and 2 (B) are shown separately.

**Figure 4.**
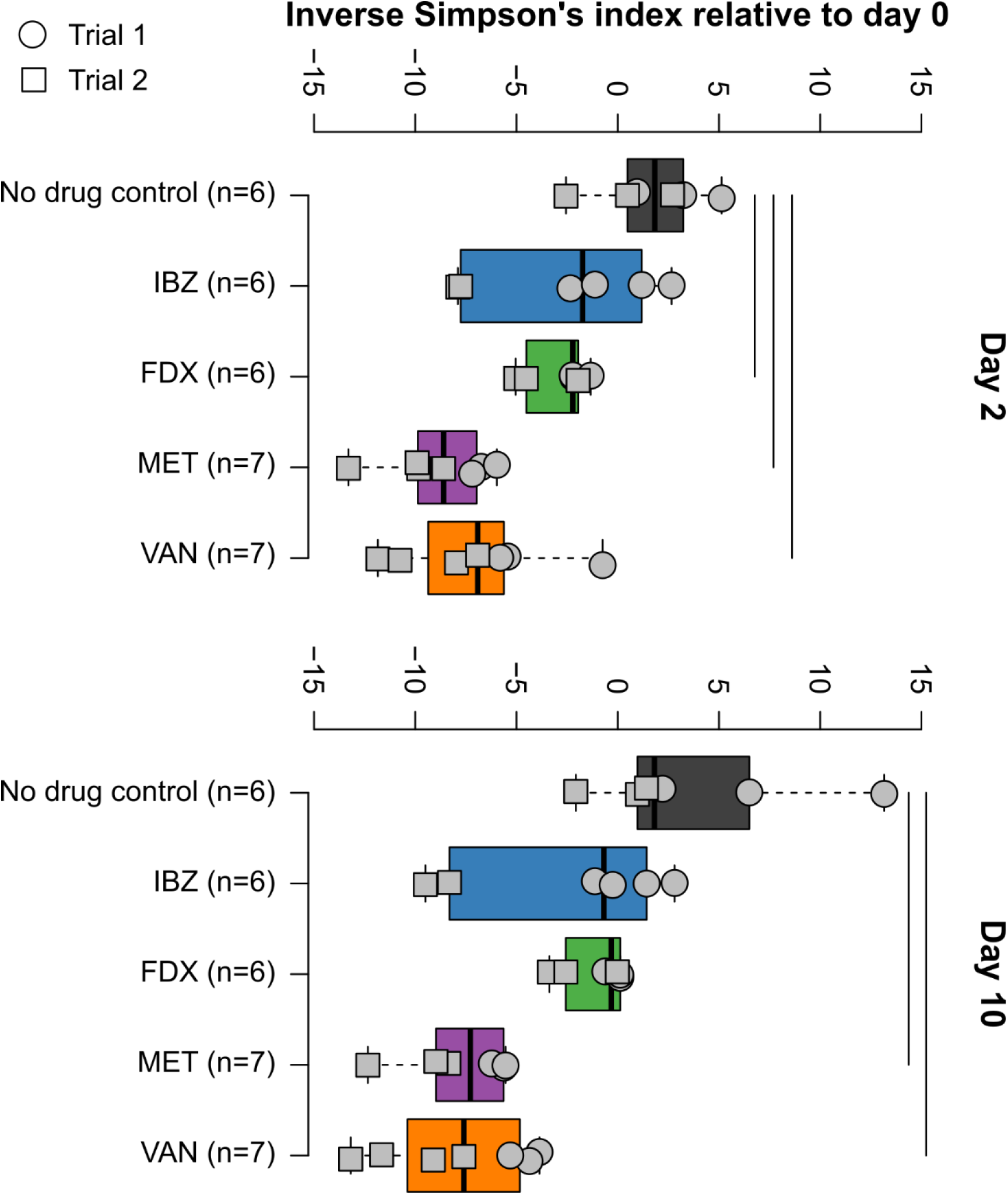
Alpha diversity (Inverse Simpson’s index) for each treatment group on days 0 (top) and 10 (bottom) of treatment. Samples from individual mice were colored gray and shown according to trial (circle = trial 1, box = trial 2). Vertical lines represent statistically different (p<0.05) group comparisons after adjustment for false discovery rate.

Linear Discriminant Analysis Effect Size (LEfSe) analyses evaluated the impact of MET, VAN, FDX, and IBZ (Figure 5 A-D) on the relative abundance of taxa at baseline (day 0) versus end of treatment (EOT, day 10). MET-treated mice exhibited the most extensive taxonomic changes (Figure 5A). Notably, taxa with higher relative abundances at EOT compared to baseline primarily belonged to the Bacillota phylum (Figure 5A). VAN-treated mice demonstrated increased relative abundances of *Acidaminococcaceae, Acidaminococcales*, and *Burkholderiales* at EOT (Figure 5B). In FDX-treated mice, a higher relative abundance of *Phocaeicola, Bacteroidaceae, Bacteroidales, Erysipelotrichaceae*, and *Erysipelotrichales* was observed at EOT (Figure 5C). IBZ-treated mice showed increased relative abundance of *Odoribacter, Odoribacteraceae*, and *Bacteroidales* at EOT (Figure 5D).

**Figure 5.**
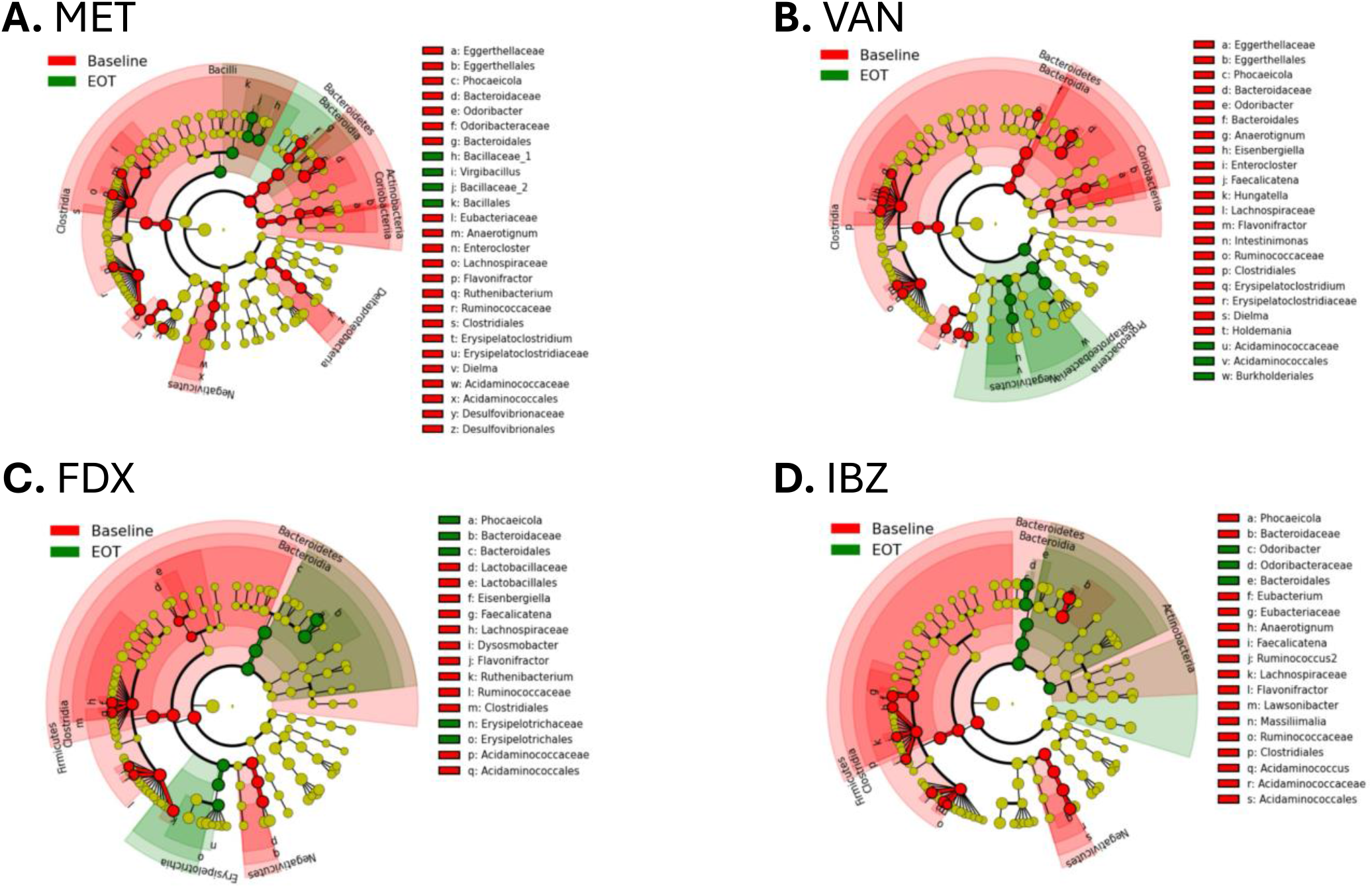
Linear discriminant analysis effect size (LEfSe) analysis evaluating the effects of: **A** Ibezapolstat (IBZ), **B** Fidaxomicin (FDX), **C** Vancomycin (VAN), and **D** Metronidazole (MET) on the relative abundance of taxa at baseline versus end of treatment (EOT). Significantly higher relative abundance taxa at the end of therapy are represented in green, while significantly higher relative abundance taxa at baseline are represented in red.

The presence-absence and relative abundance of the 15 most abundant families across both trials were evaluated (Figure 6). In trial 1, IBZ treatment significantly affected all families except *Akkermansiaceae* and *Sutterellaceae*. Additionally, seven of the 15 families exhibited significantly more reads unique to day 0, suggesting that IBZ reduced these taxa below the detection threshold. In contrast, *Odoribacteraceae* appeared to thrive with IBZ treatment, with significantly more reads unique to day 10. Similarly, shared taxa (observed on both days 0 and 10) from *Porphyromonadaceae, Desulfovibrionaceae, Rikenellaceae*, and *Odoribacteriaceae* were more abundant on day 10, suggesting they were unaffected by IBZ. Due to the limited number of mice (n=2) treated with IBZ in trial 2, no statistical analysis was performed, though the trends largely mirrored those seen in trial 1, particularly for taxa unique to day 0.

**Figure 6.**
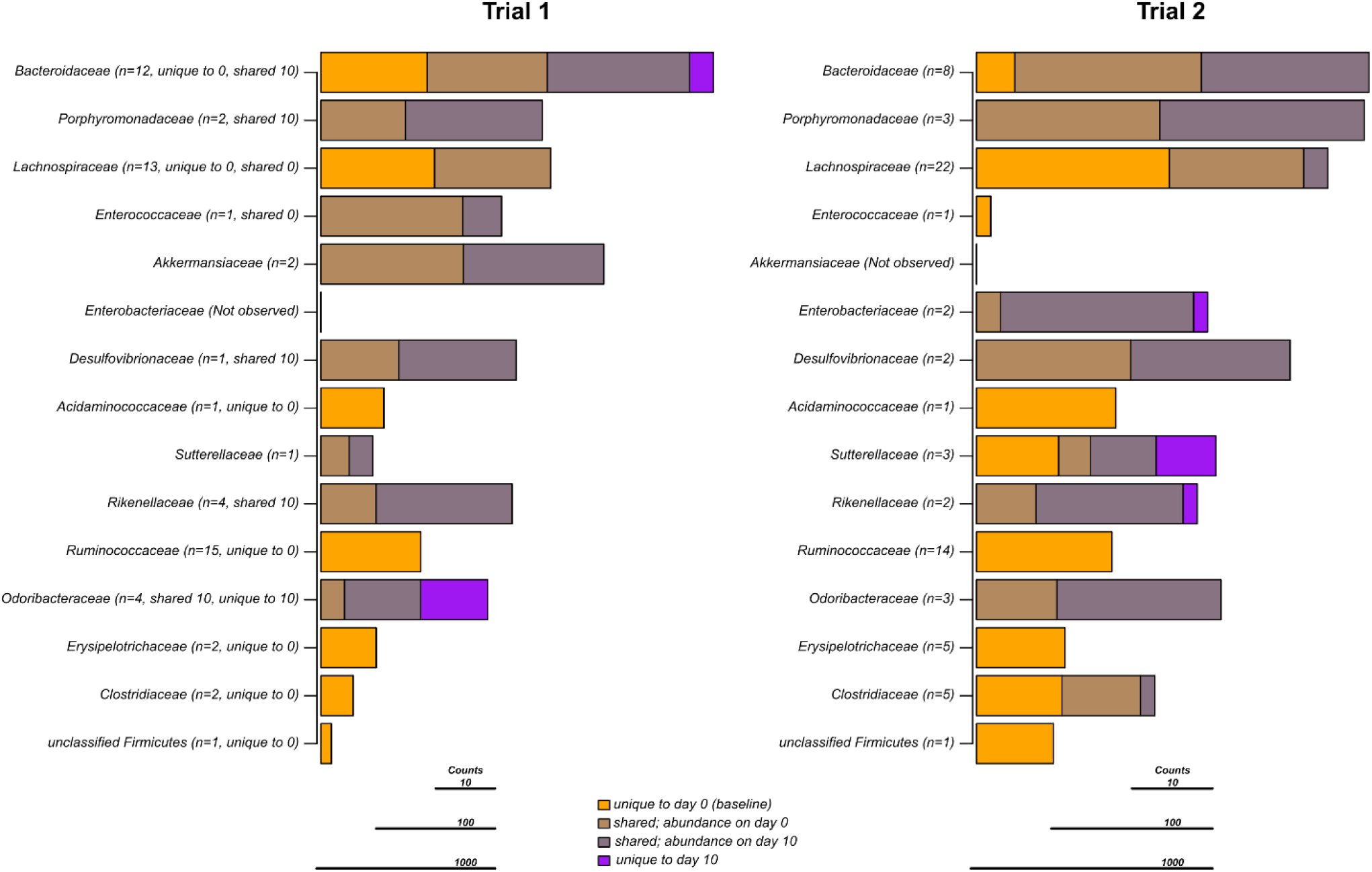
Unique and shared taxa on days 0 (baseline) and 10 (end of antibiotic exposure) for IBZ treatment. Each bar represents the median number of reads for the top 15 bacterial families observed in the study. For each family, stacked bars represent the median number of reads observed only on days 0 or 10 (i.e., unique) or observed on both days (i.e., shared). Shared bars are colored and scaled according to their relative abundance on either days 0 or 10, respectively. After each family name (in parentheses), the number of representative OTUs along with statistically significant (p<0.05) results are shown (unique to 0 = significantly more OTUs observed on day 0 compared to day 10, unique to 10 = significantly more OTUs observed on day 10 compared to day 0, shared = an overabundance of reads on day 0 or 10 as indicated).

## Discussion

### FDX, IBZ exhibits lower microbiome disruption compared to VAN or MET

IBZ is a competitive inhibitor of the Pol IIIC DNA synthesis cognate substrate 2’-deoxyguanosine 5’-triphospate (dGTP) and has completed Phase 2 studies for the treatment of CDI in adults ^10,13^. Results from the Phase I healthy volunteer study demonstrated distinct alpha and beta diversity between healthy volunteers given one of two doses of IBZ compared to those given oral VAN. This was characterized by increased abundance of Actinomycetota in IBZ-treated subjects and increased Pseudomonadota in VAN-treated subjects. In the Phase 2a study, CDI patients on IBZ therapy exhibited increased alpha diversity, marked by the regrowth of beneficial Bacillota with an increase in secondary bile acid concentrations ^9^. Most clinical trial development in CDI includes VAN as a comparator and the microbiome changes of IBZ compared to other CDI-directed antibiotics were not done during the clinical trial development process. To provide key insights into microbiome changes of IBZ compared to other CDI-directed antibiotics including FDX, we used a humanized mouse model (GF mice receiving a human-derived FMT) to assess the microbiome changes associated with IBZ, FDX, MET, and VAN. Our results demonstrated that IBZ-treated mice showed alpha diversity changes comparable to FDX, with much less alteration than those treated with MET or VAN (Figure 4). These differences in alpha diversity resulted in distinct beta diversity changes between treatment groups. Overall, IBZ and FDX had smaller beta diversity shifts from baseline compared to VAN or MET (Figure 2 and Figure 3). The taxonomic impacts of IBZ and FDX also differed, with varying regrowth patterns observed at the Phylum, Class, Order, and Family levels (Figure 5 and Figure 6). The application of multiple analysis methods provided insights beyond the use of single method. For example, the unique-shared analysis (Figure 5) indicated that IBZ had no impact on Porphyromonadaceae, Desulfovibrionaceae, Rikenellaceae, and Odoribacteriaceae, while LEfSe analysis (Figure 6) did not show these results.

Taken together, our analysis places IBZ in a similar category of microbiome disruption as FDX, indicating a narrower spectrum of microbiome alteration compared to broader-spectrum agents like VAN and MET. This opens up opportunities for future studies to differentiate these antibiotics based on their distinct effects on gut microbial composition changes. In the Phase 2a trial, *Ruminococcaceae* (Bacillota phylum) was positively correlated with increased secondary bile acid concentrations, which has an inhibitory effect on *C. difficile* spore germination ^13,23^.

### Humanized mouse models can discern antibiotic effects on the microbiome

In this study, we demonstrated complex and dynamic changes in the gut microbiome of GF mice following human-derived FMT in response to different antibiotic exposures. The model was sensitive to microbiome changes over time, even in control arms, and was also affected by dietary changes. This highlights the significant role of donor strains and laboratory conditions in shaping the trajectory of the microbiome ^24,25^. Using human-derived FMT, the post-FMT bacterial composition in mice resembled that of the human gut microbiome, primarily consisting of Bacillota and Bacteroidota phyla ^26^. However, minor differences, such as the increased abundance of Verrucomicrobiota phylum in one trial, emphasize the need for sufficient sample sizes to account for baseline variability between trials. Despite these variations, the mouse model produced results consistent with previous human trials, allowing for direct comparisons between treatments that would be difficult to achieve in human clinical trials. Characterizing both the compositional and metabolomic changes of the gut microbiome is a vital aspect in understanding its functional roles and will be the focus of future work.

Our study has several limitations. We chose to humanize the mice with healthy human microbiota to replicate the Phase I, healthy volunteer trial. In future developments of this model, we will aim to investigate microbiome-related changes in infection models, including CDI. We observed metagenomic changes consistent with published human data on IBZ and other anti-CDI agents. While this study focused on metagenomic changes in this study, future research will expand to include metabolomic and other functional analyses.

Using a humanized mouse model, we demonstrated that IBZ has a similarly narrow spectrum of activity to FDX as opposed to the broad-spectrum effect of VAN and MET. Significant differences were observed between the metagenomic data for IBZ and FDX, which may allow for further differentiation of these two narrow-spectrum antibiotics in future studies. This study also highlights the utility of humanized mouse models for evaluating the impact of antibiotics on the gut microbiome, closely mimicking the known effects in humans.

## Acknowledgments

We want to thank the staff at the Animal Resource Center (ARC) at Montana State University for their efforts with the animal experiments used in this study.

## Data Availability

All 16S rDNA sequencing reads were deposited in the National Center for Biotechnology Information (NCBI) BioProject database with accession code PRJNA934954. The authors declare that all other data supporting the study’s findings are available within this publication and its supplementary information files or from the corresponding authors upon request.

